# A Joint Distribution Approach on Non-linear Functional Connectivity

**DOI:** 10.1101/2022.07.16.500288

**Authors:** Huzheng Yang, Shi Gu

## Abstract

This report summarizes experiments on exploring non-linear functional connectivity. Using resting-state functional MRI data from the HCP1200 dataset, we define nodes as 17 functional networks and edges as the joint distribution between times series pairs. Linear dependence is removed before taking the joint distribution. We then employ a test for normality on the joint distribution to find non-normal distribution patterns. However, the result from an experimental run of 10 subjects shows that: only less than 1% of edges is non-normal distributed, and the location of such edges is not consistent across subjects. From this point of view, the non-linear part seems to be governed by random noise.

## Introduction

Classic functional connectivity uses Pearson’s R between time series to define edges, but Pearson’s R only explores linear dependence, the non-linear part remains unexplored. The goal of this study is to find patterns in the non-linear part.

Recent studies of dynamic functional connectivity (Preti et al., 2017) employ a sliding window approach to explore changes in functional connectivity over time. In theory, such changes can be modeled as non-linear interactions, but modeling such non-linear interactions is a challenging problem due to the noisiness and arbitrary timing of fMRI. Without loss of generality, we take a joint distribution approach to explore nonlinear interactions.

## Methods and Experiments

### Dataset setup

In this study, we use publicly shared data from Human Connectome Project, HCP1200 (Essen et al., 2012). In the full HCP1200 dataset, there are around 1000 subjects with 4 runs of resting-state fMRI scans, but we only use the first 10 subjects and the first run. Each run contains 900 frames for 16 minutes (TR=1000ms). The fMRI data were pre-processed using the public shared release pipeline and we downloaded the un-parcellated volumetric data.

### Network Nodes

We applied Schaefer2018 atlas (Schaefer et al., 2017) to extract ROI time series from the volumetric fMRI data. Standard Z-score normalization is then separately applied to every ROI of every subject. Then we grouped ROIs by 17 networks (Fig1). These 17 networks are used as nodes in the following analysis.

**Figure 1:**
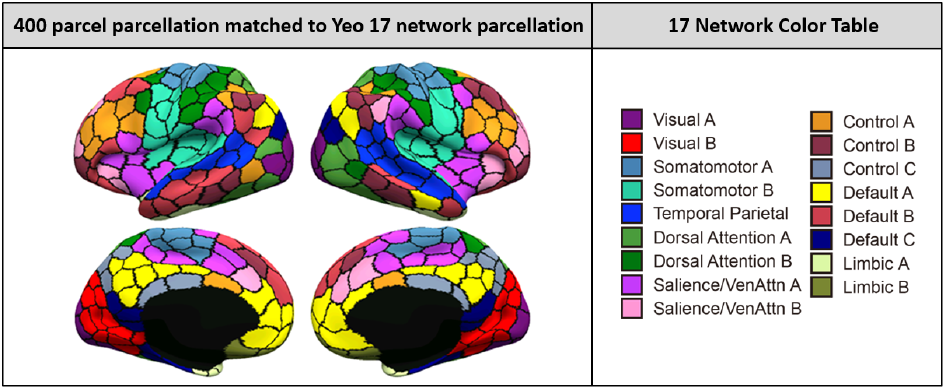
Schaefer2018 brain parcellation and 17 functional brain networks. (image taken from Schaefer2018)

### Network Edges

For every pair of nodes *v*_1_ and *v*_2_, *v*_1_ contains *N*_1_ time series from each ROI, and *v*_2_ contains *N*_2_ time series, instead of averaging time series in the same node, we take a cartesian product and treat every pair of time series as a data point, totaling *N*_1_×*N*_2_ data points. For every data point, time series pair *t*_*i*_ and *t* _*j*_, we first remove linear dependence by 2 linear regressions

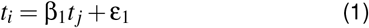

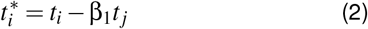

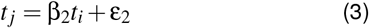

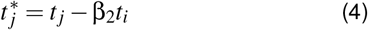

β is scalar linear regression coefficient and ε is the remaining non-linear part. We use 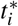 and 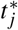 as the new time series pair. Then every frame in 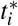 and 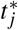 pair is treated as a data point in the joint distribution, one joint distribution contains *N*_1_ ×*N*_2_ ×*F* data points, where *N*_1_ and *N*_2_ depends on the nodes and *F* = 900 is the number of frames.

### Test for Normality

We aim to find non-normal distributed edges. For every edge, we randomly sample 10000 data points from the joint distribution and generate another 10000 data points from the 2D Gaussian distribution. Then a two-sample Two-dimensional Kolmogorov-Smirnov test (Press & Teukolsky, 1988) is applied to retrieve P-value.

The full procedure code is listed in Listing 1.

Listing 1: example code

~~~
**for** subject_i **in** [ 10 subject s ] :
 **for** node_1 **in** [ 17 networks ] :
  **for** node_2 **in** [ 17 networks ] :
     create_new_joint_distribution () ;
    **for** t_i **in** [ node_1 rois ] :
     **for** t_j **in** [ node_2 rois ] :
        remove_linear_part (t_i, t_j) ;
        add_to_joint_distribution () ;
   normality_est () ;
~~~

### Results

Table 1 shows the resulting P-values and thresholds with Bonferroni correction, 10 subject totalling 10×(17×17−17) = 2720 edges, but only 17 edges (0.7%) are non-normal distributed. Also, these non-normal distributed edges are distributed in various locations and are not consistent between subjects. Figure 2a shows some examples of non-normal distributed edges and Figure 2b shows normally distributed edges, it’s hard to tell the difference between them and it’s unclear why these edges failed to pass the test for normality.

**Table 1:** Results for normality tests.

| Total Edges | P< 0.05 | P< 0.05/<br>(17*17-17) | P< 0.05/<br>(10*(17*17-17)) |
| --- | --- | --- | --- |
| 2720 | 1396(51.3%) | 73(2.7%) | **17(0.7%)** |

**Figure 2:**
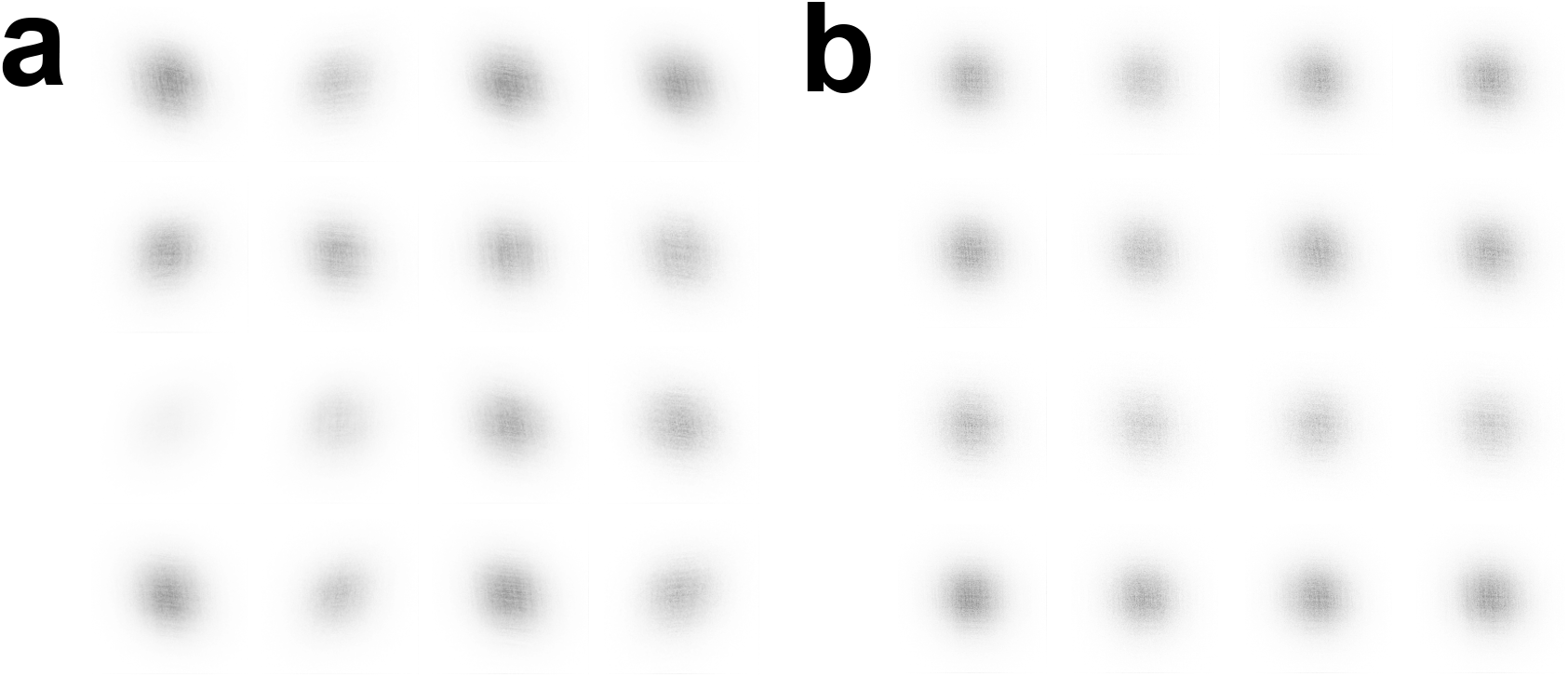
Example joint distributions. **a:** non-normal distributed. **b:** normal distributed

## Conclusion & Discussion

By using a joint distribution approach, we found the nonlinear part of functional connectivity to be governed by random noise. This may be because joint distribution throws away all the timing information, future work may design a model that accompanies timing information, but such a model also needs to address the arbitrary timing nature of fMRI and is still an outstanding challenge.

